# Atkinsonella hypoxylon virus capsid structure highlights the diversity of capsid proteins among the *Partitiviridae*

**DOI:** 10.1101/2025.08.29.673116

**Authors:** Micol Venturi, Matthew Calthorpe-Byrne, Beate Aftret, Donna McNeale, Bernd Rehm, Frank Sainsbury

## Abstract

Atkinsonella hypoxylon virus (AhV) is a fungi-infecting betapartitivirus and the typical member of the *Partitiviridae*, a family of persistent viruses that infects a broad range of organisms. Partitiviruses have been largely overlooked following their designation as cryptic viruses. However, evidence is accumulating that they play an important role in the ecology of their hosts. Since the capsid proteins of partitiviruses have been implicated in virus-host interactions, exploring their structural biology may give clues into the evolution, horizontal transmission, and host adaptation of partitiviruses. The capsid of AhV shares the same organisation of 60 dimeric capsid protein protomers arranged with T=1 icosahedral symmetry as other partitiviruses. The structure, determined by cryo-electron microscopy to 2.4 Å, shows that AhV has a unique iteration on the protrusion domain with an extensive network of hydrophobic interactions among equivalent interdigitating loops at the dimerization interface. AhV also shares a conserved helical core in the shell domain, which we extend to all genera of the recognised partitiviruses using protein structure prediction. The helical core appears to be a conserved element of the picobirnavirus lineage of capsid protein folds and provides a template onto which various elaborations of the protrusion domain have evolved. The involvement of the protrusion in virus-host interactions has previously been proposed and our findings provide evidence of a structural device enabling capsid protein diversification during the evolution of the *Partitiviridae*.

## Intro

*Partitiviridae* is a family of persistent viruses, with species now identified from fungal, plant, protist, animal and bacterial hosts. The five genera recognised by the International Committee on the Taxonomy of Viruses (Vainio et al., 2018) include the *Alphapartitivirus* and *Betapartitivirus* that infect both plants and fungi, and *Deltapartitivirus, Gammapartitivirus* and *Cryspovirus* that are restricted to plant, fungal and protist hosts, respectively. However, cryspovirus-like partitiviruses have since been found in yeast (Taggart et al., 2023) and two proposed genera, *Epsilonpartitivirus* (Nerva, Silvestri, et al., 2017) and *Zetapartitivirus* (Gilbert et al., 2019; Jiang et al., 2019), contain viruses that infect arthropods and fungi. Recently identified prokaryotic partitiviruses form a distinct clade (Le Lay et al., 2023; Neri et al., 2022; Urayama et al., 2024), leading to the recent establishment of a sister family, *Soropartitiviridae* (Turner et al., 2025). There is an emerging recognition that partitiviruses and partitivirus-like viral entities are enormously successful. In addition to the regular discovery of new hosts, they may be the most abundant viruses in fungi (Gilbert et al., 2019) and wild plants (Roossinck, 2012).

Formerly known as cryptic viruses due to the apparent asymptomatic nature of partitivirus infections, details are emerging of virus-host interactions that are beginning to shed light on the nature of their persistent infections. For example, a mutualistic relationship has been identified between pepper cryptic virus 1 and its plant host *Capsicum annum* (Safari et al., 2019) and a conditionally mutualistic relationship may exist for white clover cryptic virus 1 and its leguminous host (Nakatsukasa-Akune et al., 2005). Conversely, several fungal partitiviruses influence the virulence of their hosts (Guo et al., 2024; Xiao et al., 2014; Zheng et al., 2014), which may also be host-specific (Hyder et al., 2013). The majority of partitiviruses have two genomic dsRNAs, each containing a single open reading frame. RNA-1 encodes an RNA-dependent RNA polymerase (RdRp) and RNA-2 encodes the capsid protein (CP). Despite such a limited repertoire of viral proteins, the determinants of virus-host interactions remain largely unknown. However, at least one alphapartitivirus CP modulates host defences (Nakatsukasa-Akune et al., 2005) and structural features of the gammapartitiviruses and deltapartitiviruses have led to speculation that the CP plays a role (Byrne et al., 2021; Ochoa et al., 2008).

The isometric capsid of partitiviruses is composed of 60 quasi-symmetrical dimers arranged with T=1 icosahedral symmetry, however, there is considerable variability in the organisation of CP domains between partitivirus genera. Protrusions emanate from a conserved shell domain of deltapartitiviruses and gammapartitiviruses at distinctly different positions; at the CP dimerisation interface in deltapartitiviruses forming an intertwined spike (Byrne et al., 2021), and distal to the dimerisation interface in gammapartitiviruses forming an arch (Pan et al., 2009; Tang, Pan, et al., 2010). High-resolution structures are only available for three virus capsids from these two genera, and as a result, the structural diversity within and between partitivirus genera is not well understood. Greater understanding of partitivirus capsid structures across the family may help elucidate a possible role of the CP in adapting to persistent lifestyles.

Here we present the high-resolution capsid structure of the typical member of the *Partitiviridae*, the betapartitivirus Atkinsonella hypoxylon virus (AhV). The CP model confirms another unique iteration on the core shell-forming design principle seen in the resolved partitivirus CPs with a filled “butte” structure characterised by extensive hydrophobic interactions between reciprocal penetrating loops at the dimer interface. Combined with *in silico* prediction of CP structures from recognised genera of the *Partitiviridae*, the findings highlight the flexibility of an evolutionarily conserved core CP structure to accommodate insertions and rearrangements to drive structural divergence.

## Methods

### Capsid expression and purification

The CP coding sequence from AhV RNA-2 (NC_003471.1) was ordered as a clonal gene with flanking *Age*I and *Xho*I restriction sites in pUC57 (www.geneuniversal.com). Using these restriction sites, the CP sequence was subcloned into pEAQ-*HT* (GenBank accession GQ497234; (Sainsbury et al., 2009)). Recombinant plasmid constructs were verified with Sanger sequencing by the Griffith University DNA Sequencing Facility. The resulting pEAQ-HT-AhV_CP expression vector was maintained in *Agrobacterium tumefaciens* strain LBA4404 transformed by electroporation and propagated at 28 °C in the presence of 50 μg/mL of kanamycin, 100 µg/mL streptomycin and 50 µg/mL rifampicin. LBA4404 cultures were resuspended in infiltration buffer (10 mM MES (pH 5.6) with 10 mM MgCl2 and 100 μM acetosyringone) to an OD_600_ of 0.4 and incubated for 2-4 h at ambient temperature. Leaves of 5-week-old *Nicotiana benthamiana* plants were vacuum infiltrated using a desiccator/vacuum chamber and incubated for 6-7 days. Approximately 25 g of infiltrated tissue was extracted in 50 mL extraction buffer (50 mM MOPS (pH 7.0) with 140 mM NaCl, 0.5 mM dithiothreitol added fresh) with complete EDTA-free protease inhibitor cocktail (www.sigmaaldrich.com). The cell lysate was filtered using miracloth to remove debris and centrifuged at 25,000 g for 25 min at 4 °C. A 50% iodixanol solution was prepared using a 6x extraction buffer (120 mM MOPS (pH 7) with 840 mM NaCl), from which a cushion of 30% iodixanol in extraction buffer (50 mM MOPS (pH 7.0) with 140 mM NaCl) was used to sediment AhV VLPs at 150,000 g for 3 h at 18 °C. The cushion containing the sample was collected and diluted in extraction buffer then centrifuged at 150,000 g for 3 h at 18 °C. The pellet was resuspended with extraction buffer in a small volume (100 µL – 200 µL) by vigorously pipetting up and down.

### Negative stain TEM

Capsid preparations (4 μL, ∼0.25 mg/mL) were applied to formvar/carbon coated copper grids (ProSciTech) for 2 minutes. Grids were then washed on a droplet of water for 30 seconds two times, then stained on a droplet of Uranyless EM stain (ProSciTech) for 2 minutes. Excess stain was wicked away with filter paper and air-dried prior to storage. Images were taken on a Hitachi 7700 transmission electron microscope at 80 kV.

### Mass spectroscopy

Mass spectroscopy analysis was performed by the Australian Proteome Analysis Facility. In-gel trypsin digest was performed following reduction and alkylation of the sample resolved by SDS-PAGE. Peptides were subjected to LC-MS/MS analysis (Thermo, Ultimate 3000 HPLC coupled with Q-Exactive HFXMS) and the raw data file was processed using Proteome Discoverer (Version 2.5.0.400, Thermo Scientific). The data was searched using search engines SequestHT specifying trypsin as the enzyme with a maximum of 2 missed cleavages, precursor mass tolerance of 20 ppm, fragment mass Tolerance of 0.02 Da, allowing for dynamic modification of methionine (oxidation) and N-terminal Acetylation, and static cysteine carbamidomethylation. False discovery rates were set to <1% for peptide-spectrum matches and proteins using a custom database containing 77,048 *Nicotiana* sequences.

### Cryo-EM image processing

Image processing was carried out in RELION 4.0 (Kimanius et al., 2021). Motion correction was performed with MOTIONCOR2 (Zheng et al., 2017). The contrast transfer function of motion corrected micrographs was determined with gCTF (Zhang, 2016). Approximately 110, 000 particles were picked with the Laplacian picking tool in RELION. Particles were extracted in a 512 x 512 pixel box. The particle stack was reduced to 27, 000 through iterative 2D and 3D classification. For 3D refinement, an initial model was generated from a small subset of this stack. 3D reconstruction resulted in a model of 3.0 Å. Further CTF refinement and Bayesian polishing resulted in a final 3D reconstruction with a resolution of 2.4 Å. Detailed parameters for data collection are shown in Table 1.

**Table 1.** Cryo-EM data collection, refinement, and validation statistics.

| Data collection and processing |  |
| --- | --- |
| Sample applications to grid | 1 |
| Magnification | 81,000 |
| Voltage (kV) | 300 |
| Electron exposure (e<sup>-</sup>/Å<sup>2</sup>) | 50 |
| Defocus range of micrographs (μm) | -0.5 to -2.0 |
| Pixel size (Å) | 0.829 |
| Symmetry imposed | I |
| Initial particle images (no.) | 110, 000 |
| Final particle images (no.) | 27, 000 |
| Map resolution (Å) | 2.4 |
| FSC threshold | 0.143 |
| Number of frames | 50 |

| Refinement |  |
| --- | --- |
| Map sharpening $B$ factor ($\text{\AA}^2$) | −94.4 |
| Model composition |  |
| Protein residues | 1027 |
| Nucleic acids | 0 |
| R.m.s. deviations |  |
| Bond lengths ($\text{\AA}$) | 0.003 |
| Bond angles ($^\circ$) | 0.668 |
| Validation |  |
| Clashscore | 14.73 |
| Poor rotamers (%) | 5.07 |
| Ramachandran plot |  |
| Favoured (%) | 96.32 |
| Allowed (%) | 3.59 |
| Disallowed (%) | 0.09 |

The initial atomic model for AhV CP was built using ModelAngelo (Jamali et al., 2024) and manually modified with COOT (Emsley & Cowtan, 2004). Refinement was carried out with Phenix real space refine (Adams et al., 2010) and the atomic model was validated using MolProbity (Davis et al., 2007). All figures depicting AhV structures were prepared with UCSF Chimera and UCSF ChimeraX (Pettersen et al., 2021). AhV CP coordinates are deposited in the Protein Data Bank (8PHH) and the cryo-EM reconstruction of the capsid is deposited in the EM Data Bank (EMD-17662).

### Phylogenetic analysis and in silico structural comparisons

Sequence alignments were made using the MAFFT plugin and the L-INS-i algorithm in GeneiousPrime 2019.2.3 (www.geneious.com). A maximum likelihood tree was built using IQ-Tree 2.3.5 (Minh et al., 2020) and automated model selection using ModelFinder (Kalyaanamoorthy et al., 2017) provided by Galaxy Australia (The Galaxy, 2024). Columns where at least 40% of the sequences had gaps were masked for tree building and branch support was determined using UFBoot2 (Hoang et al., 2018). Additional cryspovirus CP sequences were identified by position-specific iterated BLAST (PSI-BLAST; blast.ncbi.nlm.nih.gov/Blast.cgi) with cryptosporidium parvum virus 1 CP (YP_009508066.1) as the query sequence. Additional deltapartitivirus sequences were selected to represent the diversity within this genus from previously identified long and short form CP sequences (Byrne et al., 2021).

The structures of 43 of 45 classified partitivirus CPs and 19 additional CPs from putative deltapartitiviruses and cryspoviruses were predicted using AlphaFold (version 2.1.2), also *via* Galaxy Australia. The highest-ranking model was selected for further analysis in all cases. Structural alignments and molecular visualisation were performed using UCSF ChimeraX (Pettersen et al., 2021).

## Results and Discussion

### Transient plant-based expression of AhV coat proteins yields assembled virus-like particles

The 652 amino acid coding sequence for AhV CP was inserted into the pEAQ-*HT* expression vector (Sainsbury et al., 2009). Following infiltrations of *A. tumefaciens* into leaves of mature *N. benthamiana* plants, expression from this vector is driven by the cauliflower mosaic virus 35S promoter and engineered cowpea mosaic virus RNA-2 untranslated regions, supported by co-expression of the P19 suppressor of post-transcriptional gene silencing from tomato bushy stunt virus. Clarified lysates of homogenised leaf tissue were applied to an iodixanol cushion to isolate AhV capsids from host cell proteins, followed by centrifugal sedimentation to remove excess iodixanol and concentrate the capsids. SDS-PAGE showed a major band at the expected ∼74 kDa (Figure 1A), and the identity of this protein was confirmed as the AhV CP by mass spectroscopy of tryptic fragments (Figure 1C; Supplementary Figure S1). Despite the apparent full-length protein on the gel, no peptides were identified from the N-terminal 80 amino acids. We attribute this to the suitability of trypsin to digest the CP, although proteolytic modifications to the N-terminus during expression cannot be ruled out. The assembly of AhV CP into virus-like particles ∼35 nm in diameter was confirmed by negative stain electron microscopy (Figure 1B). These results show that transient expression in plants is a viable approach to the recombinant expression of fungal virus-like particles. Although similar outcomes have been found for various eukaryotic viruses such as those of plants (Byrne et al., 2019; Hesketh et al., 2018), insects (Castells-Graells et al., 2021), fish (Marsian et al., 2019) and mammals (Marsian et al., 2017), it is notable that here we have not optimised the sequence for cross-kingdom heterologous expression. In addition, we have developed a purification protocol for recombinant virus capsids that isolates assembled particles from host cell components in the first step, allowing simultaneous purification, buffer exchange and concentration with a second high-speed pelleting step. The resulting capsid preparations are, as we show below, suitable for high-resolution structure determination.

**Figure 1.**
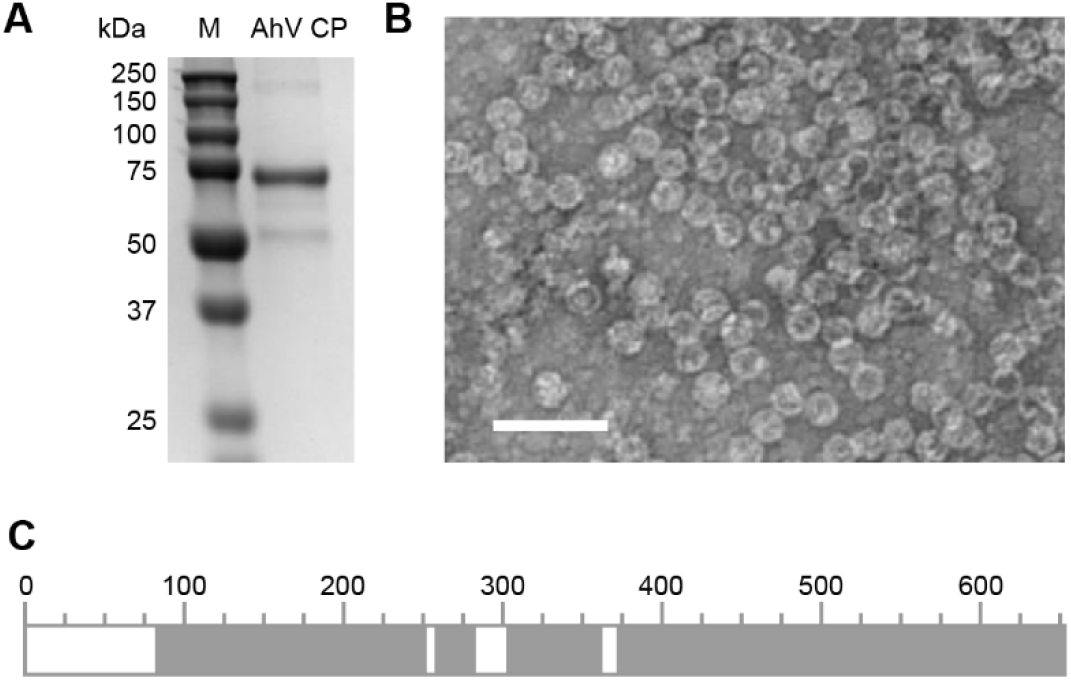
Expression and purification of AhV capsids. **(A)** SDS-PAGE gel of purified AhV showing the expected AhV CP band at ∼74 kDa. The minor band at ∼55 kDa is the large subunit of Rubisco. **(B)** Electron micrograph of purified AhV capsids. Bar = 100 nm. **(C)** Schematic representation of peptide mass fingerprinting showing the coverage (grey) of identified peptides following tryptic digest of the ∼74 kDa band in (A). Numbering refers to the AhV CP amino acid position and the sequence coverage is shown in detail in Supplementary Figure S1.

### Determination of the AhV capsid structure reveals a unique CP domain architecture

The overall capsid structure of isometric AhV capsid is similar to the available structures of the picobirnavirus lineage of dsRNA viruses (Abrescia et al., 2012), including all partitiviruses characterised to date. The capsid is composed of 120 coat proteins arranged in T = 1 symmetry, where each asymmetric unit comprises a homodimer of the AhV CP (Figure 2). Structure refinement with icosahedral symmetry imposed yielded a density map with a global resolution of 2.4 Å (Figure 2; Supplementary Figure S2). The final atomic model was refined in the presence of surrounding chains to satisfy inter-chain interactions and revealed subtle differences between the individual chains of the asymmetric unit. The model for chain A includes 566 residues (71-268, 275-590, 601-652) and chain B includes 562 residues (69-267, 274-537, 542-589, 601-651). The N-termini of both chains consists of a long disordered region that has been seen in other partitiviruses. For both chains, short loops that are missing in the model occur on the surface of the capsid and are presumably too flexible to reliably model. Superposition of the individual chains yields an RMSD of 0.4 Å (Supplementary Figure S3). Variations in backbone can be seen for short loops near the 5-fold and 3-fold symmetry axes of the capsid, as would be anticipated for quasi-symmetric monomers.

**Figure 2.**
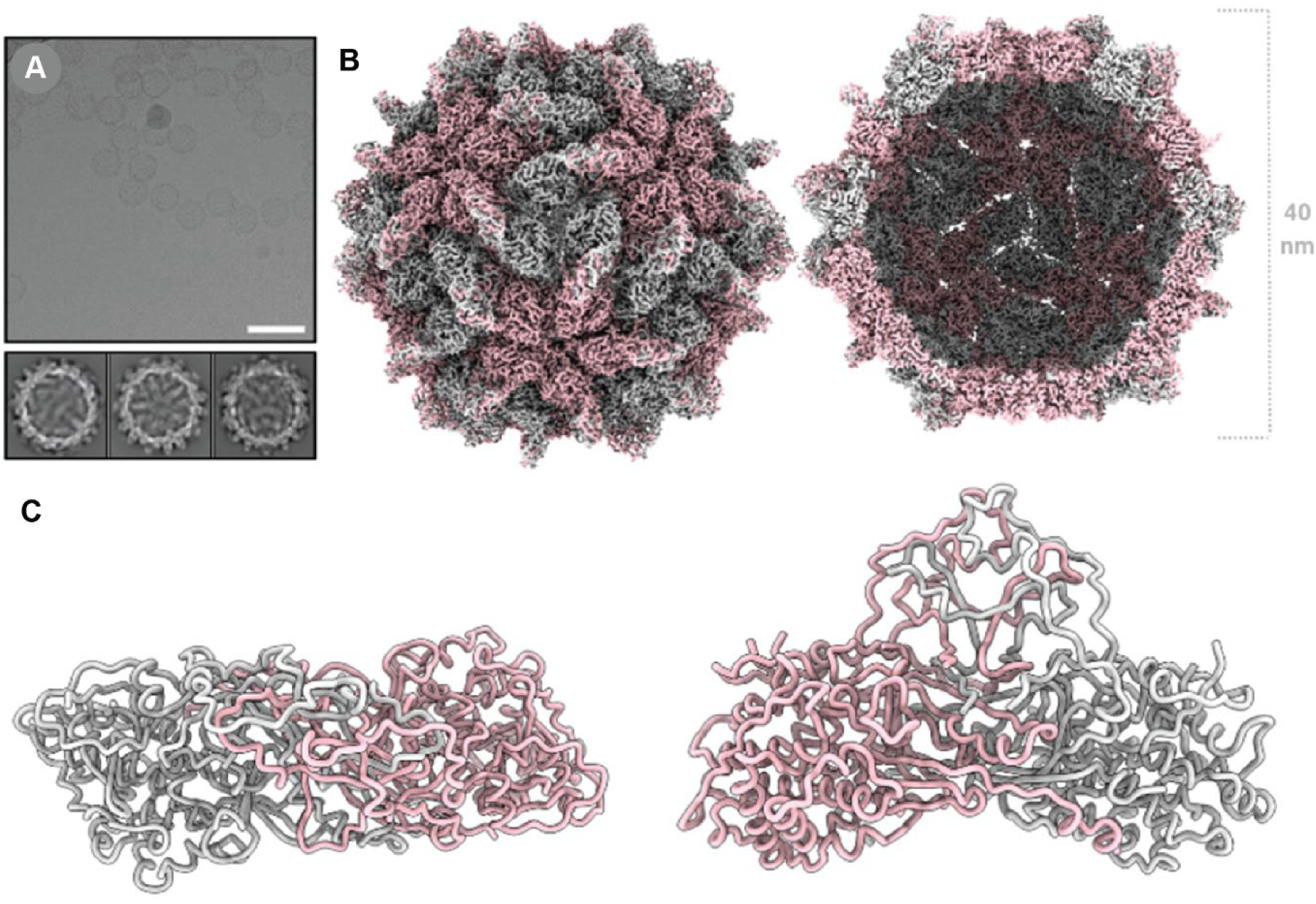
AhV capsid structure. **(A**) A section of a representative micrograph of AhV VLPs and 2D class averages. Scale bar = 100 nm. **(B)** The 2.4 Å resolution 3D reconstruction of AhV, coloured according to quasi-conformer, with monomer A coloured pink and monomer B coloured grey. The capsid surface (left) and a central slice through the capsid (right) are shown. **(C)** The atomic model of AhV, showing the model for a single asymmetric unit from the bottom (left) and the side (right) with the monomers coloured as in panel B.

The AhV CP forms a T=1 icosahedron of 60 dimeric protomers comprised of two distinct domains: the shell (S) and protrusion (P) domains. The S domain (res. 71-409) adopts a predominantly α- helical fold with seven α-helices (α1-α7) arranged around a central, extended 24-residue helix (α3, res. 125-148). This arrangement creates a roughly rhomboidal shape that is characteristic of partitivirus shell domains (Figure 2B; Supplementary Figure S4). The N-terminal helix α2 (res. 98-111) contains ARG111, which forms a stabilising salt bridge with GLU75, anchoring the N-terminal region to the underside of the shell domain of the opposing monomer. Domain swapping by the CP N-terminus is a feature that appears to be common among the partitiviruses. In betapartitiviruses, the N-terminus also contributes to the dimerization interface, with a largely hydrophobic contact patch (res. 71-85) involving ILE71, PHE72, and LEU78, complemented by the GLU75 to ARG111 salt bridge. Unlike the arch-forming P domain of the gammapartitiviruses (Pan et al., 2009; Tang, Pan, et al., 2010) and the spike-like P domain of the deltapartitiviruses (Byrne et al., 2021), the P domain of the betapartitiviruses takes the form of a butte (Figure 1C), as previously described for capsid structures at lower resolution (Tang, Ochoa, et al., 2010; Xiao et al., 2014). The high resolution of the AhV capsid reported here enables elaboration of the interactions that form the protrusion unique to the betapartitiviruses. The P domain (res. 410-652) constitutes the major dimerization interface, burying a substantial surface area of approximately 13,380 Å^2^. The domain features a modified β-sheet structure and multiple interdigitated loops. The P domain achieves dimerization through an extensive hydrophobic interface that forms the core of the dimer interaction, anchored by hydrogen bonds, such as that between GLN566 to TYR419 (Figure 3Ai). The central dimerization region contains multiple interaction hot spots, including ARG558 (24 contacts), LEU545 (22 contacts), and ASP564 (20 contacts), which create an extensive network of stabilizing interactions. The C-terminal interface region (res. 630-650) features unique interactions including a cation-π interaction between PHE633 and ARG636 and a symmetric ARG636 to ARG636 pairing between opposing monomers (Figure 3Aii). The CP dimer most likely represents the assembly unit for partitiviruses (Ochoa et al., 2008). However, these interactions are more extensive and more varied than has previously been seen in the partitiviruses. The evolution of such a complex and tight binding interface is intriguing and raises the question as to whether the dimer performs other functions in addition to its structural role in the capsid. As discussed below, this has been seen in other dsRNA viruses. In the absence of a reverse genetic system for most persistent viruses like AhV, further investigation of the functions of partitivirus proteins may be facilitated by recombinant expression approaches such as that described in this study.

**Figure 3.**
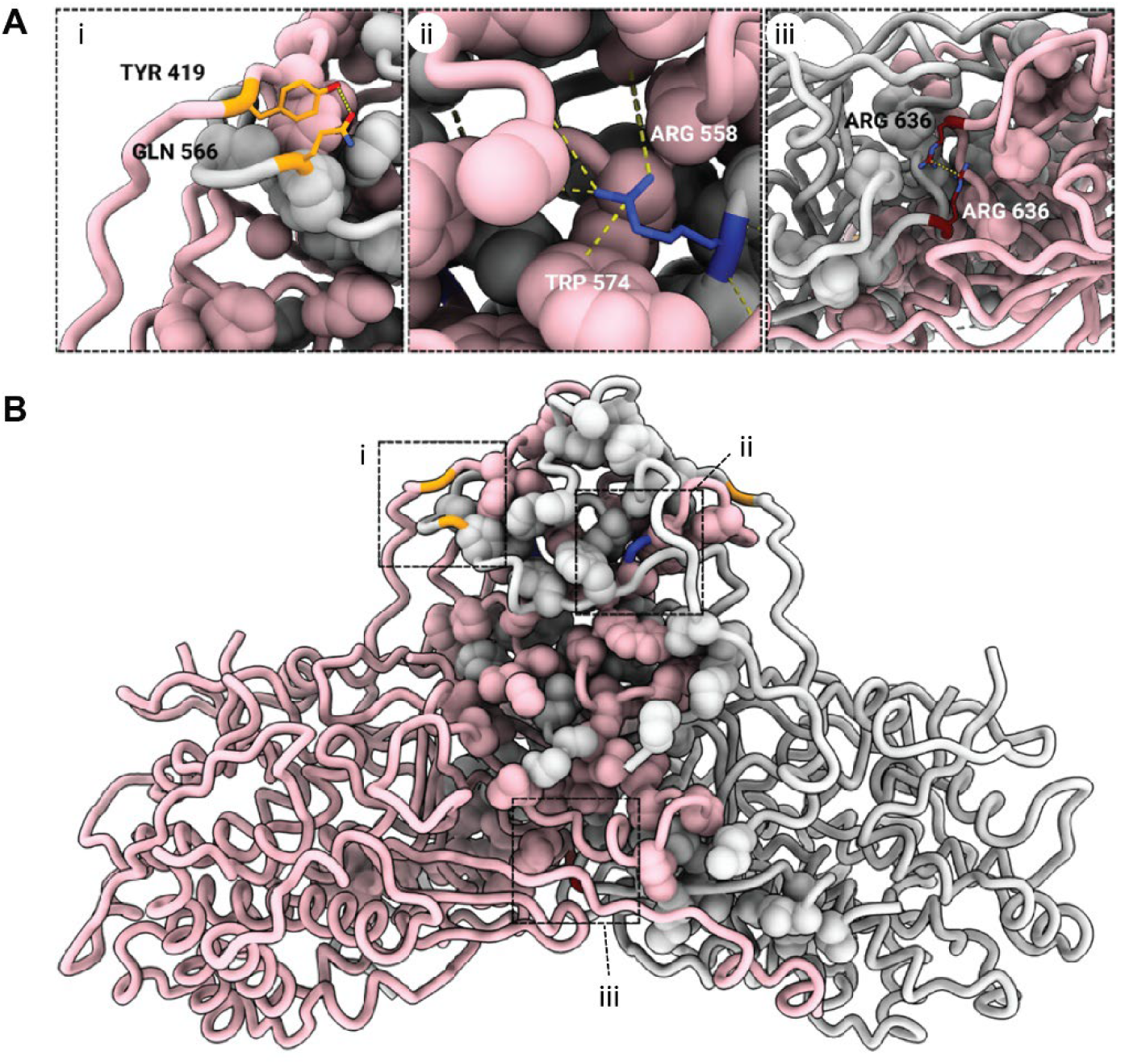
AhV capsid protein interface. **(A)** Detail expansions corresponding to dashed boxes in B showing (i) the GLN566 to TYR419 hydrogen bond anchoring an extended invading loop from the opposite monomer, (ii) the extensive contacts within the hydrophobic interface, and (iii) the symmetrical ARG636 to ARG636 pairing. **(B)** Side view of the CP dimer with the hydrophobic residues of the dimer interface shown as space-filling models. Monomer A is coloured pink and monomer B is coloured grey.

### CP structures highlight considerable structural diversity among partitiviruses

Including the AhV CP structure reported here, four partitivirus capsid structures have been resolved to high resolution: two gammapartitiviruses, and one each from the deltapartitiviruses and the betapartitiviruses. To investigate whether the remaining genera, specifically the alphapartitiviruses and cryspoviruses, have evolved yet different CP domain architectures, we used AlphaFold2 (Jumper et al., 2021) to predict the structures for an additional 61 CP sequences. To generate a broader picture for each genus, we expanded the ICTV-defined family to include 6 cryspoviruses (an increase of 5 from the 1 formally recognised member) and 20 deltapartitivuses (an increase of 15 from 5 formally recognised members), in addition to the formally recognised 17 alphapartitiviruses, 14 betapartitiviruses and 8 gammapartitiviruses (see Supplementary Tables 1-6). Using human picobirnavirus as an outgroup, phylogenetic analysis of the 65 partitivirus CP sequences grouped the CPs into clades representing the expected genera based on the published RdRp phylogeny (Vainio et al., 2018), with the exception of penicillium stoloniferum virus S, which could not be reliably grouped with the gammapartitiviruses (Figure 4A; Supplementary Figure S5).

**Figure 4.**
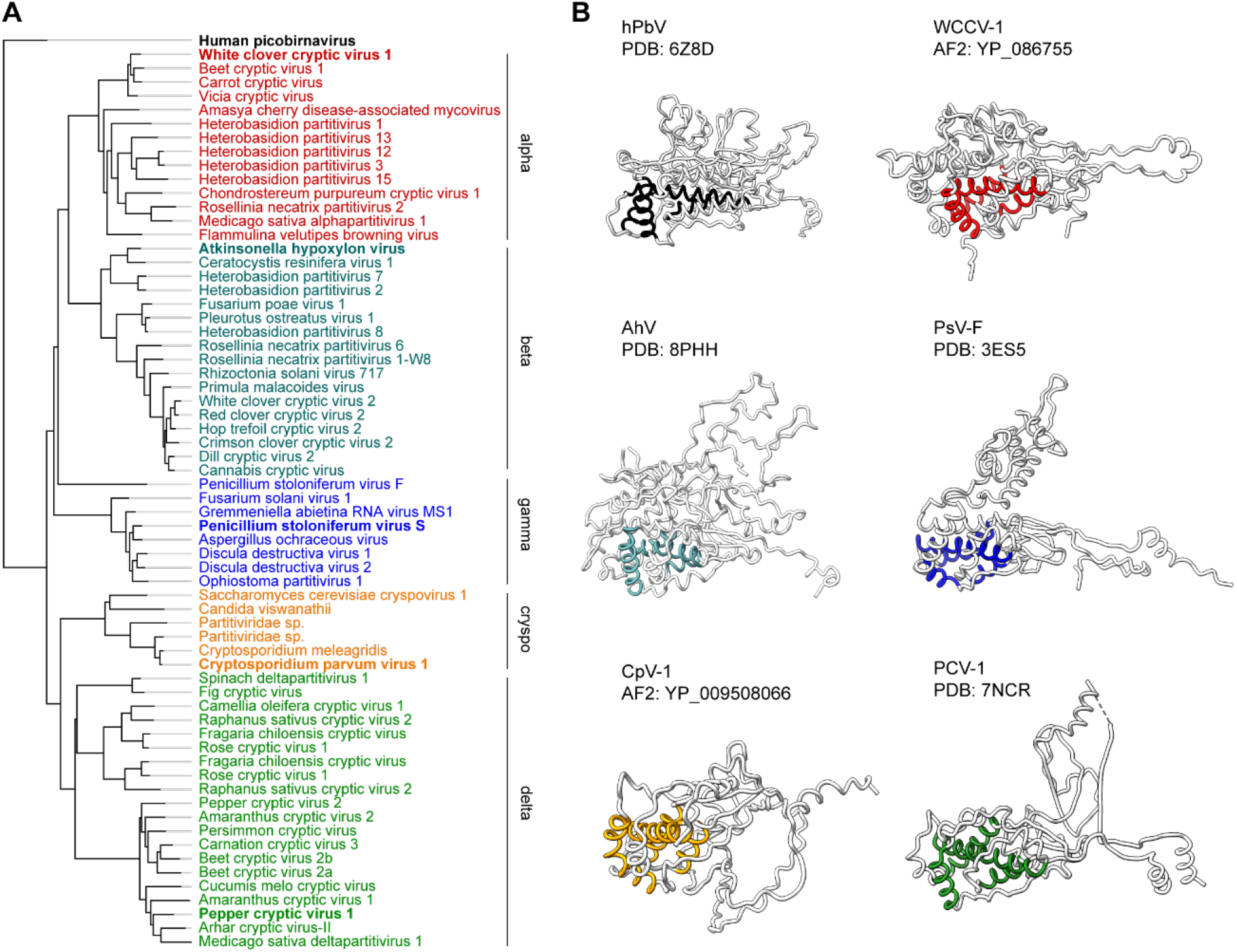
CP domain architecture in partitiviruses. **(A)** Phylogram generated using IQ-Tree (Minh et al., 2020) and the Q.pfam+F+R4 evolution model (Minh et al., 2021) with the human picobrinavirus CP (YP_239360.1) used as an outgroup. The typical member of each genus is shown in bold. A compressed tree is shown for convenience and the full tree with branch support values is presented in Supplementary Figure S5. **(B)** Molecular models of the CP monomers of representative members of each genus derived from empirically determined structures (indicated by PDB code) or predicted using AlphaFold2 (AF2; indicated by CP accession number). The colour of the conserved helical core corresponds to the genus members shown in A. hPbV is human picobirnavirus, WCCV-1 is white clover cryptic virus 1, AhV is atkinsonella hypoxylon virus, PsV-F is penicilliun stoloniferum virus F, CpV-1 is cryptosporidium parvum virus 1, and PCV-1 is pepper cryptic virus 1.

Structural alignment confirms the phylogenetic analysis of partitivirus CP sequences with highly similar structures within genera, and diverse CP architectures between them. For all CP predictions, the shell domain, particularly the core helices, were predicted with high (>90%) confidence (Supplementary Figures 6-11). Comparison of experimentally determined structures to their alphafold2 predictions shows very good agreement in backbone RMSD, and that the orientation of major structural features for the betapartitiviruses, gammapartitiviruses and deltapartitiviruses was also correctly predicted (Supplementary Figures S7, S8, and S10). It should be noted that neither the PCV-1 (7NCR) nor AhV (8PHH) structures were included in the training set of the AlphaFold version used. The predicted structures for alphapartitiviruses and cryspoviruses show that these genera also have differences in predicted CP domain architectures compared to the structurally characterised genera (Figure 4B). The relatively short cryspovirus CP (Supplementary Figure S9) sequences possess the conserved core helices and an extended N-terminus, but appear to consist of a minimal shell domain with no obvious protrusion. Alphapartitiviruses (Supplementary Figure S5) appear to have a mostly helical cap above the core helices and a long N-terminal loop that likely crosses the dimerisation interface to interact with the other monomer of the presumed protomer, as is characteristic of the dimerisation interactions within the family.

### Core helices are a highly conserved feature that accommodates structural diversity among partitiviruses CPs

Although the overall architecture of the CP is strikingly different between partitivirus genera, the shell domain of all the partitivirus CPs analysed contains a recognisable core of four α-helices (Figure 4B). In the betapartitivirus, AhV, these helices correspond to α2, α3, α6, and α12 (Supplementary Figure S4). Previously identified in gammapartitiviruses and deltapartitiviruses (Byrne et al., 2021), here we show that it is present in all genera, in addition to the Picobirnavirus CP, supporting the idea that it is a conserved element of the picobirnavirus CP structural lineage. Alignment of the core helices, hereafter termed αA-αD, shows that the 3D organisation of the core, including the direction of the helices, is highly conserved (Figure 5A). A central long helix (αB) forms the backbone of the core and it is crossed by an initial helix (αA) that runs away from the dimerisation interface to the distal edge of the protomer and a smaller helix (αC) returning back to towards the interface in the same direction as αB. A fourth helix (αD) runs perpendicular to the first three, running from the outside of the capsid to the inner surface. This organisation also holds for human picobirnavirus (Supplementary Figure S12). These data indicate that the core helices may indeed comprise the minimal fold of the shell domain of the partitiviruses (Byrne et al., 2021), although it remains to be seen whether this holds for newly discovered partitiviruses in insects and sister families in prokaryotes.

**Figure 5.**
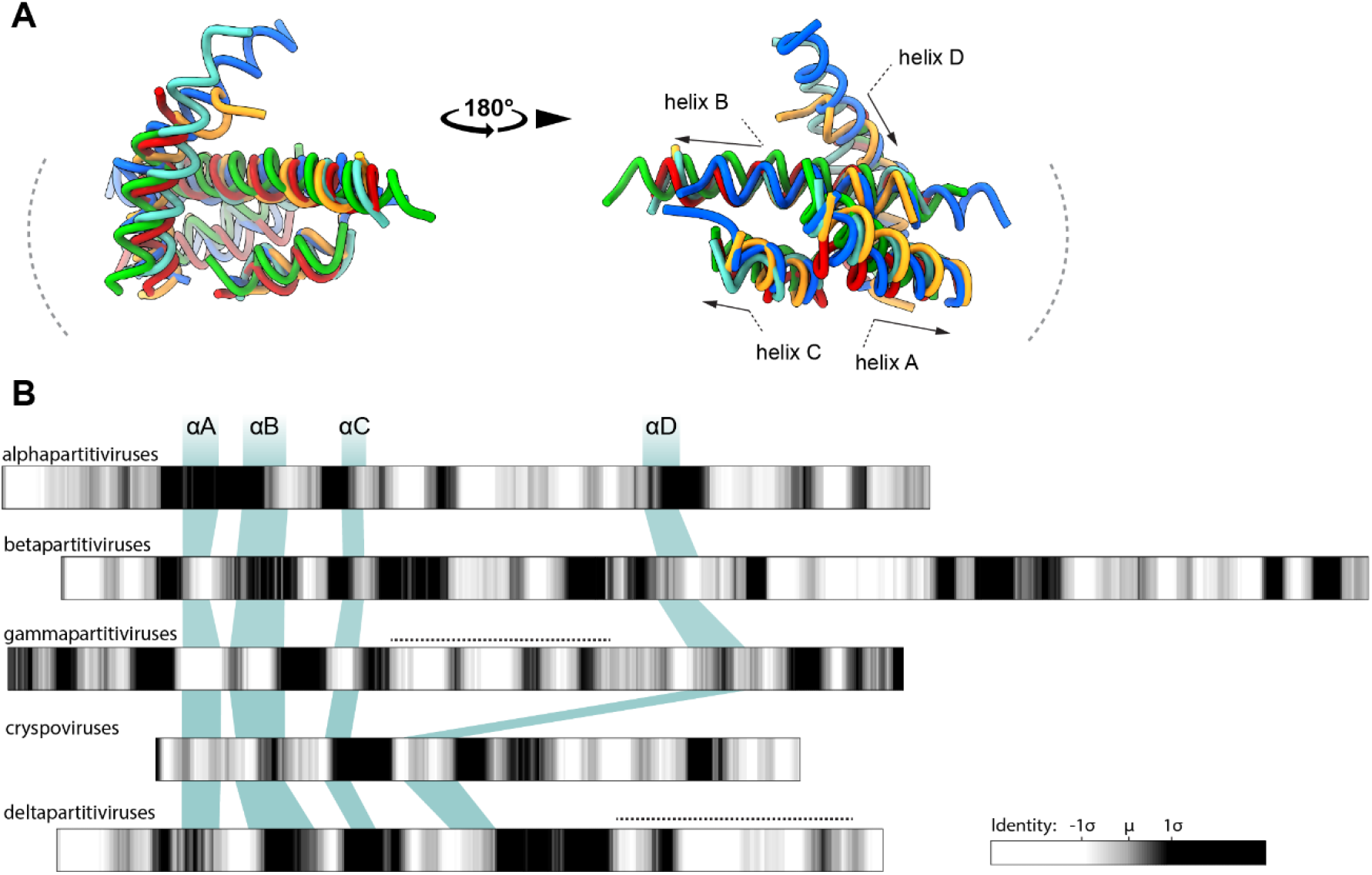
The conserved helical core of the shell domain in partitiviruses. **(A)** Alignment of the core helices of representative CP structures from each recognised partitivirus genus, annotated with the direction of the helix. The colour of the conserved helical core corresponds to the individual genus members shown in figure 4B: alphapartitivirus (AF2 - YP_086755) in red; betapartitivirus (PDB - 8PHH) in cyan; gammapartitivirus (PDB - 3ES5) in blue; cryspovirus (AF2 - YP_009508066) in yellow; and deltapartitivirus (PDB - 7NCR) in green. **(B)** Position of the core helices within the primary structure of each partitivirus genus with the intra-genus sequence identity shown as a grey-scale heat map scaled to the mean identity calculated over a 10-residue sliding window. Dotted lines represent the protrusions in the gammapartitiviruses and deltapartitiviruses.

A subset of deltapartitiviruses have putative CPs that are shorter than the demarcation criteria of the genus. No virions or capsids have been observed for the so-called short form deltapartitiviruses. However, the predicted structures of putative CPs from the short form deltapartitiviruses possess the core helices and have the same organisation and orientation as the other partitiviruses (Supplementary Figure S11). This appears to support their role as putative CPs. Moreover, some short form deltapartitiviruses appear to be tripartite, with a second putative CP. In these cases, despite sharing as little as ∼50% sequence identity, both CP-like sequences are predicted to have very similar structures (Supplementary Table S6; Supplementary Figure S11).

Despite the structural conservation of the core helices, they do not correspond to regions of sequence conservation within genera (Figure 5B). Moreover, their position in the primary sequence varies considerably between genera. In the deltapartitiviruses and cryspoviruses, the core helices are grouped together in the N-terminal half of the protein. In alphapartitiviruses, betapartitiviruses, and gammapartitiviruses αD is separated from the first three core helices by an insertion of approximately 150 amino acids (Figure 5B). In both the experimentally derived betapartitivirus structure and predicted alphapartitivirus structure, the insertion consists of an α-helical ‘cap’ over the S domain. In the gammapartitiviruses, the insertion forms the mostly helical P domain that forms an arch over the CP dimer. This is in stark contrast to the deltapartitivirus P domain at the dimerization interface and entirely C-terminal to the core helices (Figure 5B). In AhV and other betapartitiviruses, the protrusion forming the dimerization interface is also C-terminal to the final core helix αD, albeit discontinuous with elements of the S domain cap. As has been noted for other dsRNA viruses (Mata et al., 2017), it is tempting to speculate that the core helices provide an evolutionarily conserved fold that allows for the insertion of functional sequences and domains.

Viruses with persistent lifestyles and exclusively vertical transmission, such as partitiviruses, are predicted to evolve towards maximising the reproduction of their hosts (García-Ordóñez & Pagán, 2024). Evidence of mutualism in plant partitiviruses (Nakatsukasa-Akune et al., 2005; Safari et al., 2019) and ecological effects on fungal hosts (Guo et al., 2024; Hyder et al., 2013; Xiao et al., 2014) indicate host adaptation, but the viral determinants are not known. It has been shown that partitivirus RdRps are functional across host kingdoms (Nerva, Varese, et al., 2017; Telengech et al., 2024), implying that replication and transcription may not restrict host range. The CPs of dsRNA viruses are known to acquire auxiliary functions that mediate host interactions, such as cap-snatching (Fujimura & Esteban, 2011) and possibly other enzymatic activities (Mata et al., 2017). In addition, the CPs of many RNA viruses are known to regulate transcription and translation of their hosts. Therefore, although ascribing adaptive traits that enable transfer to new hosts within the RdRp cannot be ruled out, the CP of partitiviruses may play an important role.

## Conclusions

AhV is the typical member of the *Partitiviridae*. The high-resolution structure of the AhV capsid reported here confirms that the betapartitiviruses share a consistent CP and capsid organisation with other partitiviruses. It reveals an extensive hydrophobic dimerization interface that consists of multiple penetrating loops from opposite dimers, anchored by hydrogen bonds. It appears to be a highly evolved interface, including a symmetrical interaction between dimers, that reinforces the essential role of the dimer in capsid assembly. Analysis of the CPs, including structural prediction, across the recognised genera of the partitiviruses shows that there is a high-level of intra-genus structural relatedness and divergent architectures between genera. The family is characterised by a conserved shell constructed around a helical core that appears to be characteristic of the CP fold in the picobirnavirus lineage. Our results show that the shell domain, particularly the core helices, represents a structural template that the *Partitiviridae* have used to elaborate a variety of protrusions that define the genera. Unlike the genera for which high-resolution structures previously existed, the *Deltapartitivirus* and *Gammapartitivirus*, the genus *Betapartitivirus* includes members with both plant and fungal hosts. It is, therefore, interesting for the study of viruses that may have undergone horizontal virus transfer as has been suggested in partitiviruses by phylogenetic analysis. The establishment of infections in heterologous or model hosts could help shed light on the virus-host interactions that restrict or enable host range expansion. A complementary approach is structural virology, and here we have established a reference structure for this genus.

## Acknowledgements

This work was partly performed on the traditional lands of the Yugarabul, Yuggera, Jagera and Turrbal peoples. FS acknowledges support from the Commonwealth Scientific and Industrial Research Organisation (CSIRO) in the form of a Synthetic Biology Future Science Platform Fellowship. MV and DM were supported by Commonwealth Scientific and Industrial Research Organisation (CSIRO) Synbio PhD Top-up Scholarships. FS has found the time to put this manuscript together thanks to an ARC Future Fellowship (FT230100084). The work was supported by Galaxy Australia, a service provided by Australian BioCommons and its partners. This service receives NCRIS funding through Bioplatforms Australia, as well as The University of Melbourne and Queensland Government RICF funding. The authors acknowledge the facilities of Microscopy Australia at the Centre for Microscopy and Microanalysis, The University of Queensland, enabled by NCRIS. This study also used NCRIS-enabled Australian Proteome Analysis Facility (APAF) infrastructure.

