## Supplementary Figures and Tables for "Atkinsonella hypoxylon virus capsid structure highlights the diversity of capsid proteins among the *Partitiviridae*"

**Contents:**

Figure S1 – Sequence coverage by peptide mass fingerprinting

Figure S2 – Local resolution and Fourier shell correlation

Figure S3 – Superposition of chain A and chain B monomers

Figure S4 – Capsid protein secondary structure composition

Figure S5 – Phylogram of the Partitiviruses based on CP sequence

Figure S6 – *Alphapartitivirus* structural alignment

Table S1 – *Alphapartitivirus* CP sequences included in this study

Figure S7 – *Betapartitivirus* structural alignment

Table S2 – *Betapartitivirus* CP sequences included in this study

Figure S8 – *Gammapartitivirus* structural alignment

Table S3 – *Gammapartitivirus* CP sequences included in this study

Figure S9 – *Cryspovirus* structural alignment

Table S4 – *Cryspovirus* CP sequences included in this study

Figure S10 –*Deltapartitivirus* (long form CP) structural alignment

Table S5 – *Deltapartitivirus* (long form CP) sequences included in this study

Figure S11 – Structural analysis of short form CP deltapartitiviruses

Table S6 – Short form *Deltapartitivirus* CP sequences included in this study

Figure S12 – Conservation of core helices including human picobirnavirus CP

```
MSSNDSAQTRNLQEERFNERTSTPTVVTAVLPDTNGPTTNSTSGSVGPPHPTPNVPVPTQ
SSSDPPSASGIFAKEIDLPRNVIQHSGNKFILDVVPDSRFPTFAITEFVQRSFSNFTFEQ
YSYVSPASLVGYLVYMIHAFVFLVDAFERSPMSAYASEIDASHAYLRIIDAFSDAYIPDF
LFEILDTYLSHRLDIRSKLEMNVSYGSVLYKYDAPRIVAPSIFLLAHNQLISQSRESTAY
EKWLDSIVIHYSRAVIRVGNLVGGLYQSSHGSTTTHFTYRNWFARSLSRLADSATHRTHL
RRPMISEFDYNIPSVNNNTYNPYVHLLMLEPNNRNITLDFIRSLSSFCSTELKATRTLRD
HISRRSAAISRCVIKGPEAPTWHSSPLDDLKEKSKQGNFSQFCEVAKFGLPRKENSESYT
FKFPKDASTIDTAFYLIQENGRSSVLDPTTADEELHTEGMNLLFDPYDDESSAHYATVLS
GKLIQNSNIDGETLLLPDPTTGLARTNSRYLQGSVLIRNVLPEFDQHEIRLFPRYPQISR
LSASLTLLFNMRQVWIPRFKQKVDEQPKLSNFSWNEGCDGTVPSLNVVTAESSTNGPAAE
QQVILWSSYRHVSNSDRPTVDTVYYYSTLELLFGTRSSMMQTYNLHQLLSLH
```

**Figure S1. Sequence coverage by peptide mass fingerprinting.**

Coverage is indicated in red for the peptides identified following in-gel trypsin digest of the AhV CP.

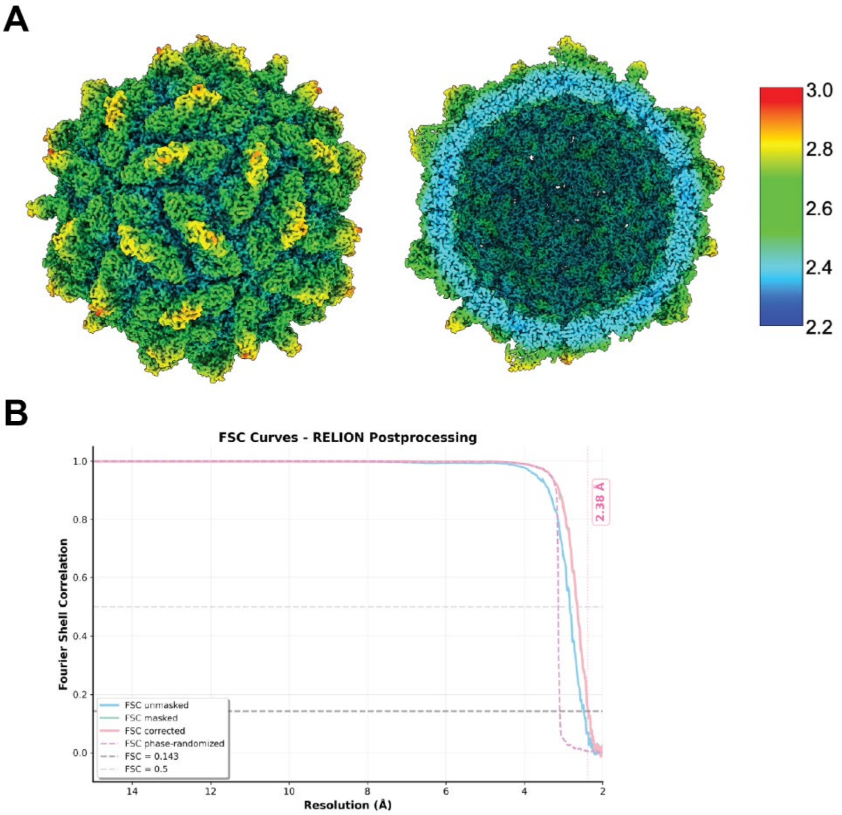


**Figure S2. Local resolution and Fourier shell correlation.**

**(A)** Local resolution filtered map of AhV VLP coloured according to resolution, showing capsid surface (left), and a central slice (right). **(B)** Fourier shell correlation – the resolution that corresponds to an FSC coefficient of 0.143 is 2.4 Å.

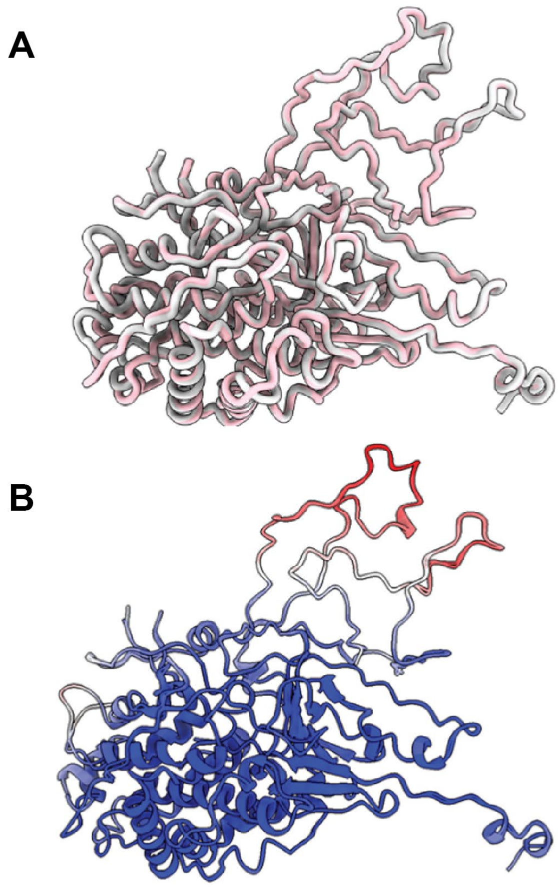


**Figure S3. Superposition of chain A and chain B monomers.**

Superposition of chain A and chain B from a single asymmetric unit of the AhV capsid. **(A)** Superposed monomers, shown as backbone ribbons for clarity. Coloured according to chain with chain A coloured pink, and chain b coloured grey. **(B)** Superposed AhV monomers, coloured according to B factor, from low (blue) to high (red).

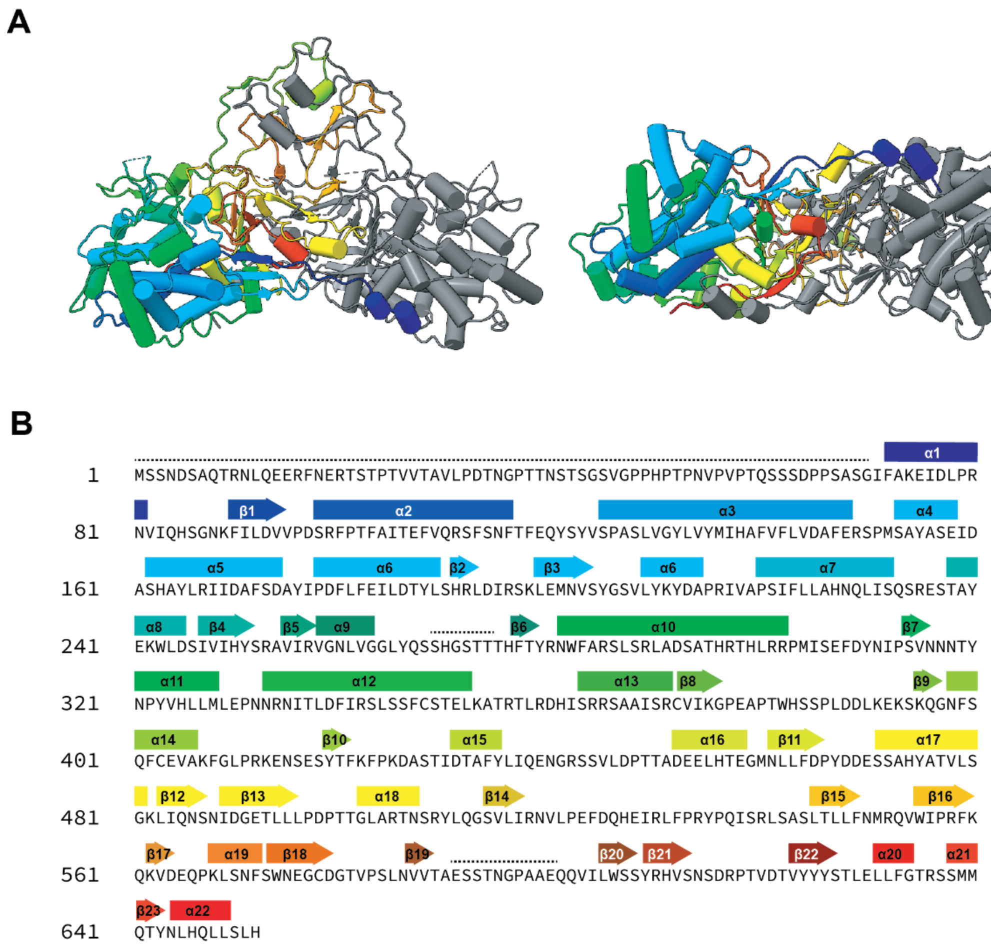


**Figure S4. Capsid protein secondary structure composition.**

**(A)** AhV CP shown in cartoon form viewed from the side (left) and the bottom (right) with monomer A coloured from N (blue) to C terminus (red). Monomer B is coloured grey for clarity. **(B)** AhV CP sequence labelled with secondary structure elements from chain A. Regions of disorder, that are not present in the final 3D reconstruction are denoted by a dashed line.

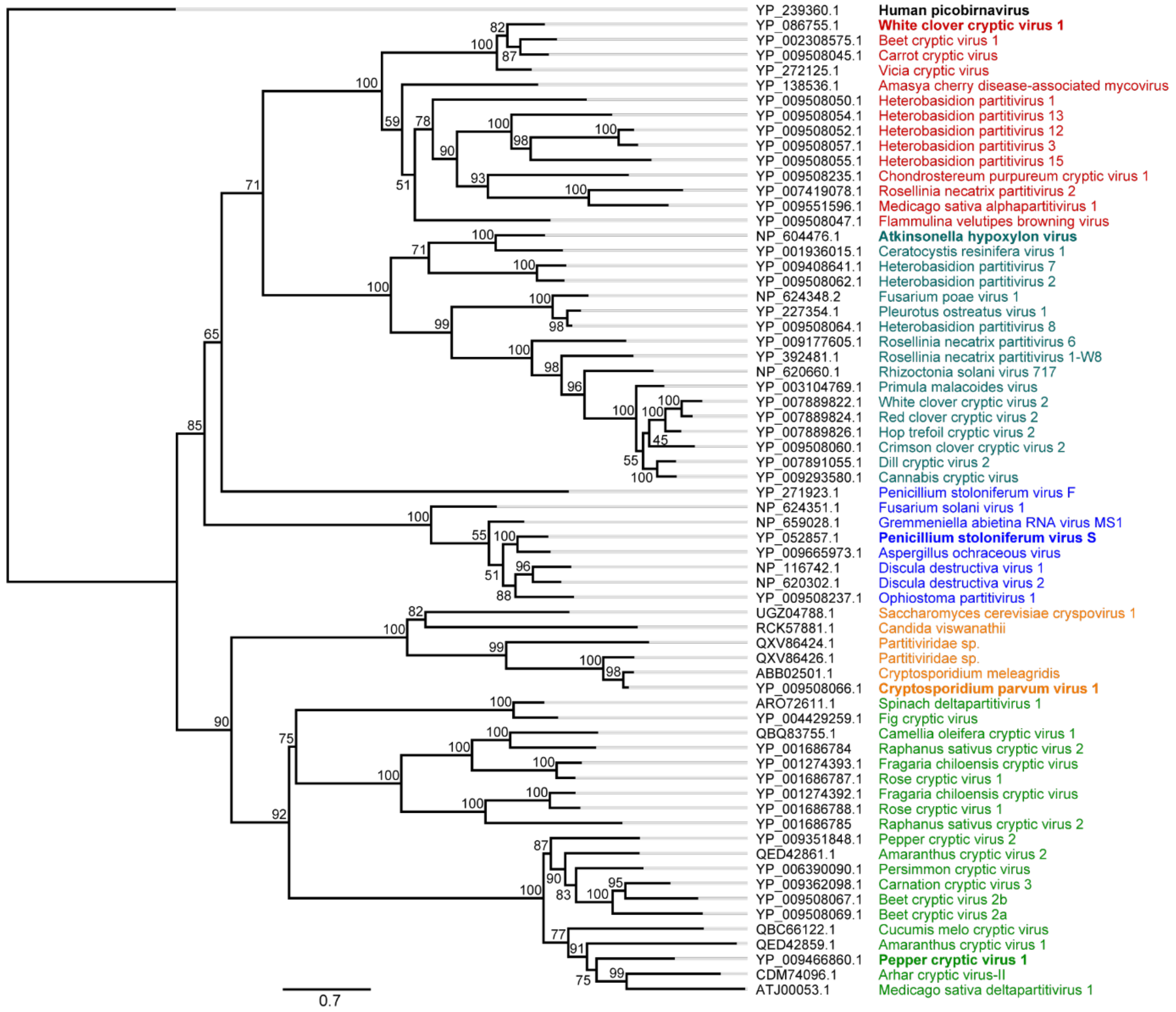


**Figure S5. Phylogram of the Partitiviruses based on CP sequence.**

The maximum likelihood tree was constructed IQ-Tree (1) and the Q.pfam+F+R4 evolution model (2). The initial alignment was created using the MAFFT algorithm L-INS-I and columns where at least 40% of the sequences had gaps were masked. Branch support is indicated at internal nodes and was determined using UFBoot2 (3) with values given as a percentage of 1000 replicates. The typical member of each genus is shown in bold.

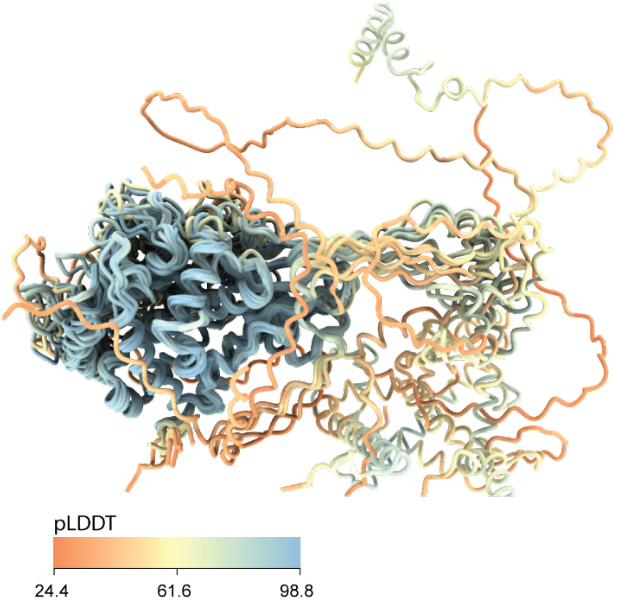


**Figure S5. *Alphapartitivirus* structural alignment.**

The predicted CP sequences of alphapartitiviruses were aligned using the matchmaker command in ChimeraX version 1.6 using white clover cryptic virus 1 as the reference. The structures include those predicted by alphafold2 (Table S1) and are coloured by the predicted local distance difference test (pLDDT) scores.

**Table S1. *Alphapartitivirus* CP sequences used in this study.**

| Accession | Virus name | Length (aa) |
|---|---|---|
| YP_009508047.1 | Flammulina velutipes browning virus | 463 |
| YP_009508235.1 | Chondrostereum purpureum cryptic virus 1 | 480 |
| YP_007419078.1 | Rosellinia necatrix partitivirus 2 | 483 |
| YP_086755.1 | White clover cryptic virus 1 | 487 |
| YP_272125.1 | Vicia cryptic virus | 487 |
| YP_002308575.1 | Beet cryptic virus 1 | 489 |
| YP_009508045.1 | Carrot cryptic virus | 490 |
| YP_009551596.1 | Medicago sativa alphapartitivirus 1 | 491 |
| YP_138536.1 | Amasya cherry disease-associated mycovirus | 504 |
| YP_009508054.1 | Heterobasidion partitivirus 13 | 509 |
| YP_009508050.1 | Heterobasidion partitivirus 1 | 510 |
| YP_009508055.1 | Heterobasidion partitivirus 15 | 512 |
| YP_009508052.1 | Heterobasidion partitivirus 12 | 520 |
| YP_009508057.1 | Heterobasidion partitivirus 3 | 521 |

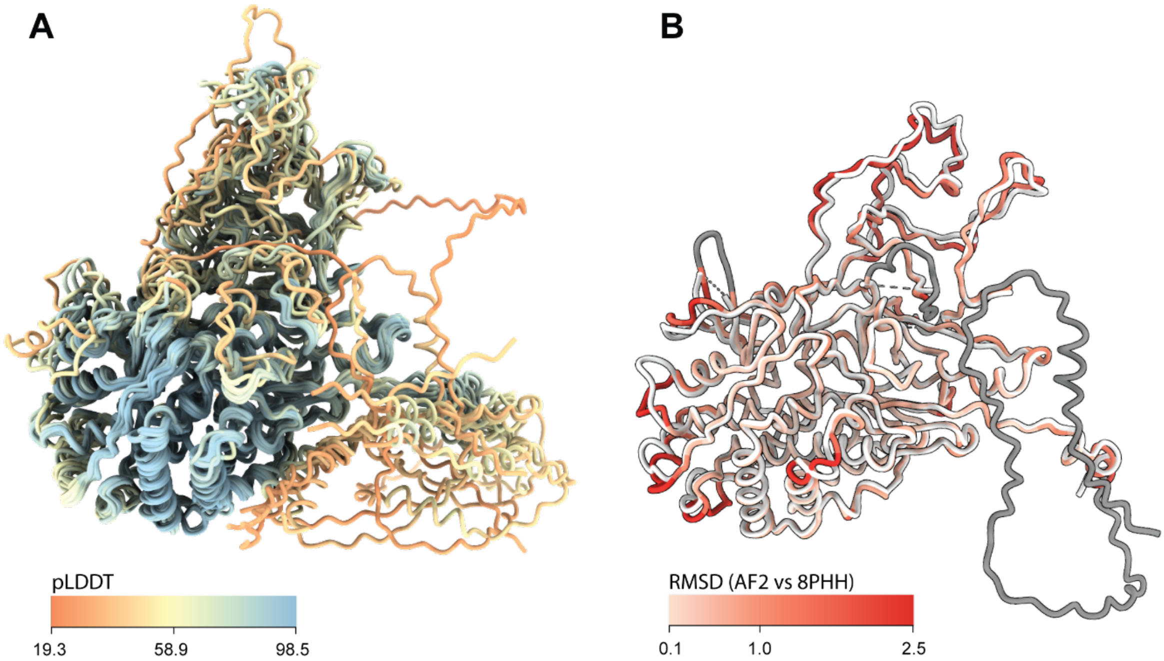


**Figure S7. *Betapartitivirus* structural alignment.**

**(A)** The predicted CP sequences of betapartitiviruses were aligned using the matchmaker command in ChimeraX version 1.6 using atkinsonella hypolxylon virus (AhV) as the reference. The structures include those predicted by alphafold2 (Table S2 below) and are coloured by the predicted local distance difference test (pLDDT) scores. **(B)** Alignment of the alphafold2 prediction and experimentally determined AhV CP structure (PDB 8PHH), coloured by root mean square deviation.

**Table S2. *Betapartitivirus* CP sequences used in this study.**

| Accession | Virus name | Length (aa) |
|---|---|---|
| YP_227354.1 | Pleurotus ostreatus virus 1 | 636 |
| NP_624348.2 | Fusarium poae virus 1 | 637 |
| YP_009508064.1 | Heterobasidion partitivirus 8 | 638 |
| NP_604476.1 | Atkinsonella hypoxylon virus | 652 |
| YP_009408641.1 | Heterobasidion partitivirus 7 | 654 |
| YP_009508062.1 | Heterobasidion partitivirus 2 | 659 |
| YP_001936015.1 | Ceratocystis resinifera virus 1 | 661 |
| YP_009293580.1 | Cannabis cryptic virus | 672 |
| YP_003104769.1 | Primula malacoides virus | 673 |
| YP_007889822.1 | White clover cryptic virus 2 | 673 |
| YP_007889824.1 | Red clover cryptic virus 2 | 673 |
| YP_007889826.1 | Hop trefoil cryptic virus 2 | 673 |
| YP_007891055.1 | Dill cryptic virus 2 | 673 |
| YP_009508060.1 | Crimson clover cryptic virus 2 | 674 |
| NP_620660.1 | Rhizoctonia solani virus 717 | 683 |
| YP_392481.1 | Rosellinia necatrix partitivirus 1-W8 | 686 |
| YP_009177605.1 | Rosellinia necatrix partitivirus 6 | 729 |

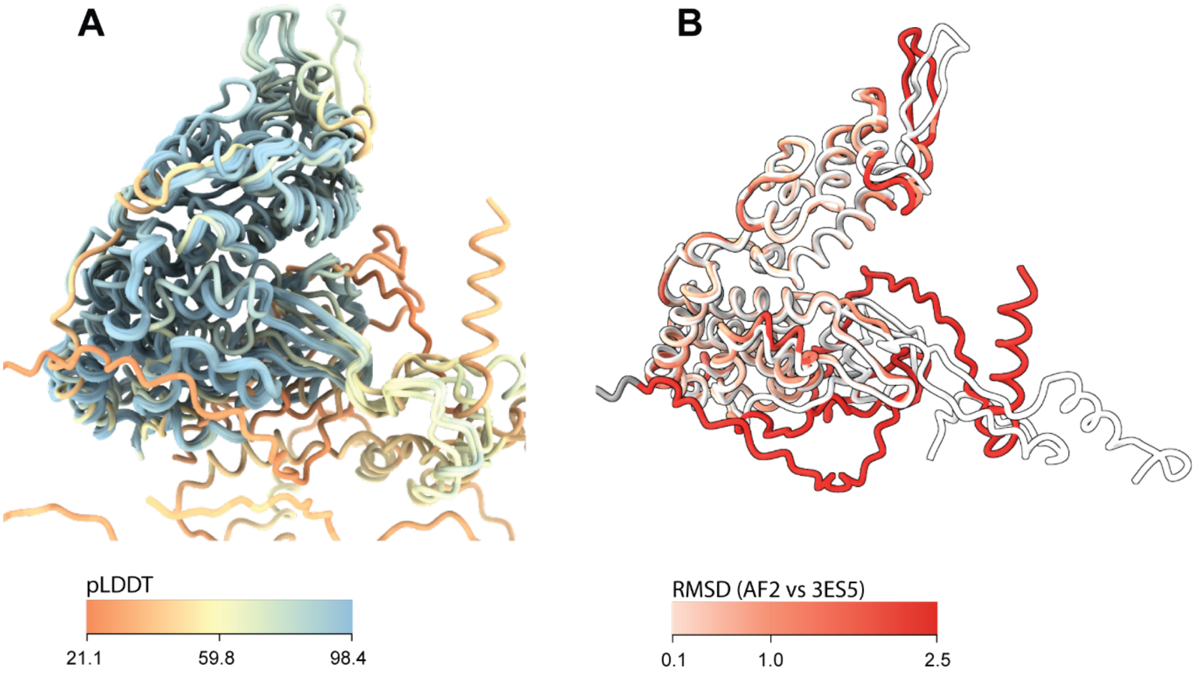


**Figure S8. *Gammapartitivirus* structural alignment.**

**(A)** The predicted CP sequences of gammapartitiviruses were aligned using the matchmaker command in ChimeraX version 1.6 using penicillium stoloniferum virus F (PsV-F) as the reference. The structures include those predicted by alphafold2 (Table S3 below) and are coloured by the predicted local distance difference test (pLDDT) scores. **(B)** Alignment of the alphafold2 prediction and experimentally determined PsV-F CP structure (PDB 3ES5), coloured by root mean square deviation.

**Table S3. *Gammapartitivirus* CP sequences used in this study.**

| Accession | Virus name | Length (aa) |
|---|---|---|
| NP_624351.1 | Fusarium solani virus 1 | 413 |
| YP_271923.1 | Penicillium stoloniferum virus F | 420 |
| YP_009508237.1 | Ophiostoma partitivirus 1 | 430 |
| YP_009665973.1 | Aspergillus ochraceous virus | 433 |
| YP_052857.1 | Penicillium stoloniferum virus S | 434 |
| NP_116742.1 | Discula destructiva virus 1 | 436 |
| NP_620302.1 | Discula destructiva virus 2 | 442 |
| NP_659028.1 | Gremmeniella abietina RNA virus MS1 | 443 |

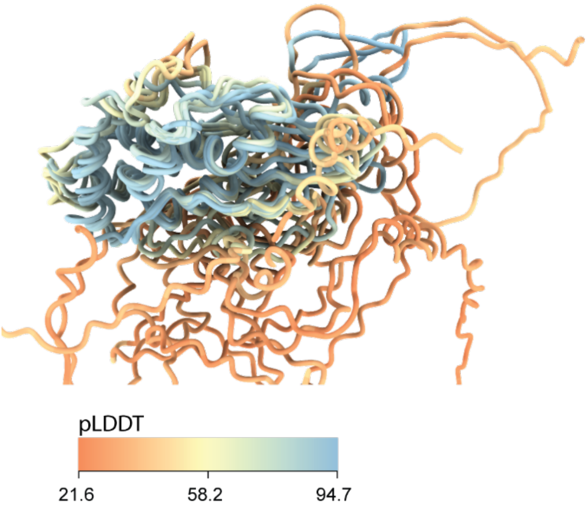


**Figure S9.** ***Cryspovirus*** **structural alignment.**

The predicted CP sequences of cryspoviruses were aligned using the matchmaker command in ChimeraX version 1.6 using cryptosporidium parvum virus 1 as the reference. The structures include those predicted by alphafold2 (Table S4 below) and are coloured by the predicted local distance difference test (pLDDT) scores.

**Table S4.** ***Cryspovirus*** **CP sequences used in this study.**

| **Accession** | **Virus name** | **Length (aa)** |
|---|---|---|
| RCK57881.1 | Candida viswanathi virus[a] | 307 |
| UGZ04788.1 | Saccharomyces cerevisiae cryspovirus 1[a] | 310 |
| ABB02501.1 | Cryptosporidium meleagridis virus[a] | 319 |
| YP_009508066.1 | Cryptosporidium parvum virus 1 | 319 |
| QXV86424.1 | Partitiviridae sp.[a] | 358 |
| QXV86426.1 | Partitiviridae sp.[a] | 366 |

a - not formally assigned to cryspoviruses

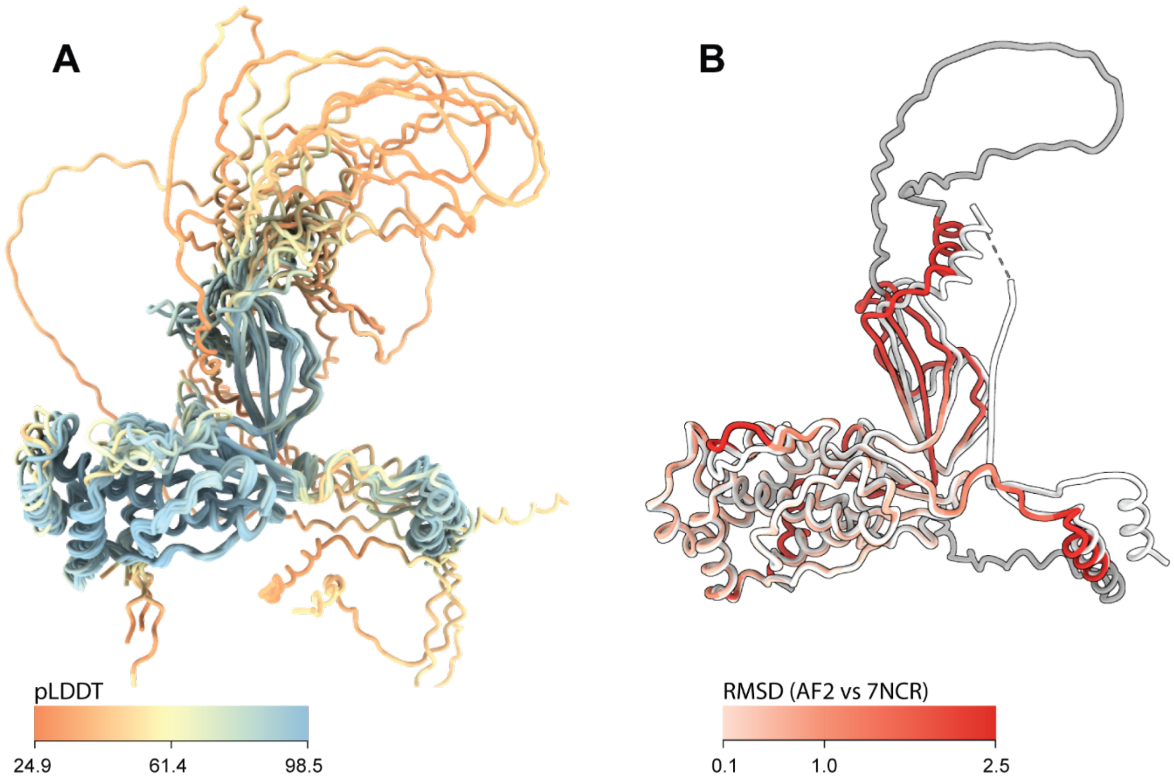


**Figure S10.** ***Deltapartitivirus*** **(long form CP) structural alignment.**

**(A)** The predicted CP sequences of long form deltapartitiviruses were aligned using the matchmaker command in ChimeraX version 1.6 using pepper cryptic virus 1 as the reference. The structures include those predicted by alphafold2 (Table S5 below) and are coloured by the predicted local distance difference test (pLDDT) scores. **(B)** Alignment of the alphafold2 prediction and experimentally determined PCV-1 CP structure (PDB 7NCR), coloured by root mean square deviation.

**Table S5.** ***Deltapartitivirus*** **(long form CP) sequences included in this study.**

| **Accession** | **Virus name (genome segment for tripartite viruses** | **Length (aa)** |
|---|---|---|
| CDM74096.1 | Arhar cryptic virus-II[a] | 389 |
| YP_009508069.1 | Beet cryptic virus 2 (RN2b) | 393 |
| ATJ00053.1 | Medicago sativa deltapartitivirus 1[a] | 394 |
| YP_009362098.1 | Carnation cryptic virus 3[a] | 411 |
| YP_009466860.1 | Pepper cryptic virus 1 | 412 |
| YP_006390090.1 | Persimmon cryptic virus[a] | 415 |
| QED42861.1 | Amaranthus cryptic virus 2[a] | 421 |
| YP_009508067.1 | Beet cryptic virus 2 (RNA2a) | 426 |
| YP_009351848.1 | Pepper cryptic virus 2 | 430 |
| QED42859.1 | Amaranthus cryptic virus 1[a] | 432 |
| QBC66122.1 | Cucumis melo cryptic virus[a] | 480 |

a - not formally assigned to deltapartitiviruses

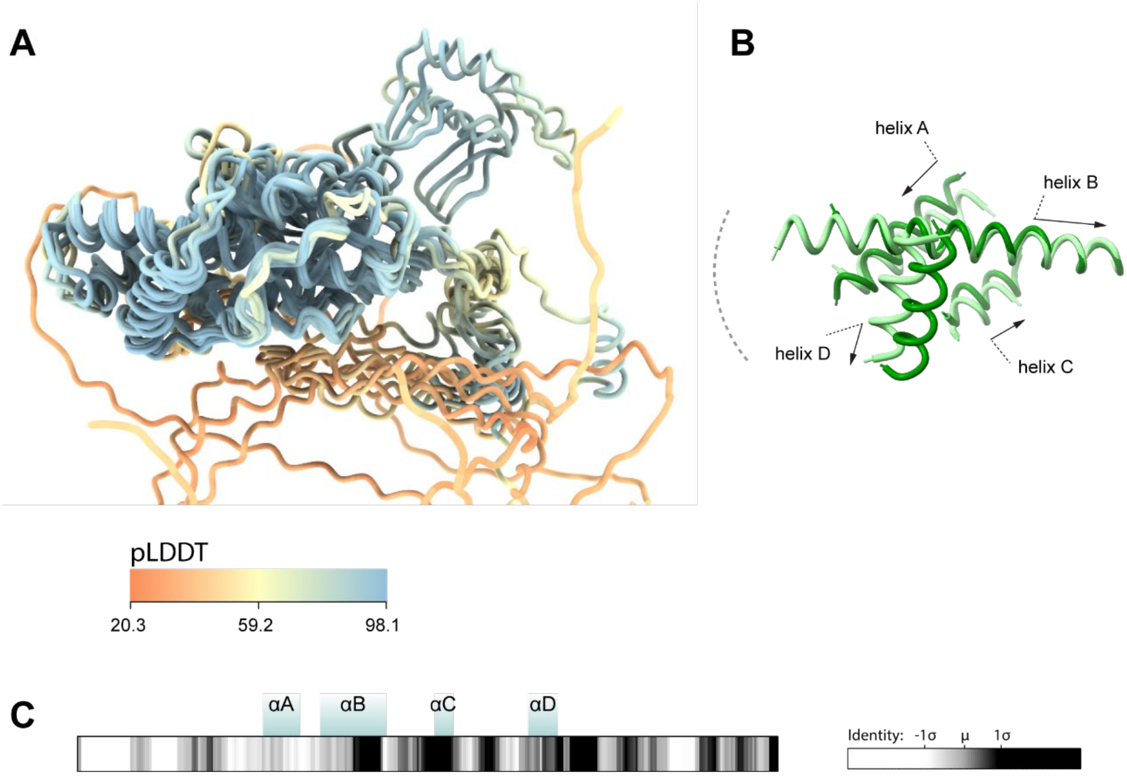


**Figure S11. Structural analysis of short form CP deltapartitiviruses.**

**(A)** The predicted CP sequences of short form deltapartitiviruses were aligned using the matchmaker command in ChimeraX version 1.6 using fig cryptic virus as the reference. The structures include those predicted by alphafold2 (Table S6 below) and are coloured by the predicted local distance difference test (pLDDT) scores. **(B)** Alignment of the core helices of the short form (light green) and long form (dark green) deltapartitiviruses, annotated with the direction of the helices. **(C)** Position of the core helices within the primary structure of the short form deltapartitiviruses with the sequence identity shown as a grey-scale heat map scaled to the mean identity calculated over a 10-residue sliding window.

**Table S6. Short form *Deltapartitivirus* CP sequences included in this study.**

| **Accession** | **Virus name (genome segment for tripartite viruses)** | **Length (aa)** |
|---|---|---|
| ARO72611.1 | Spinach deltapartitivirus 1[a] | 335 |
| YP_004429259.1 | Fig cryptic virus | 337 |
| QBQ83755.1 | Camellia oleifera cryptic virus 1[a] | 344 |
| YP_001274392.1 | Fragaria chiloensis cryptic virus[a] (RNA3) | 346 |
| YP_001686784 | Raphanus sativus cryptic virus 2[a] (RNA2) | 346 |
| YP_001686788.1 | Rose cryptic virus 1[a] (RNA3) | 346 |
| YP_001686785 | Raphanus sativus cryptic virus 2[a] (RNA3) | 347 |
| YP_001274393.1 | Fragaria chiloensis cryptic virus[a] (RNA2) | 348 |
| YP_001686787.1 | Rose cryptic virus 1[a] (RNA2) | 348 |

a - not formally assigned to deltapartitiviruses

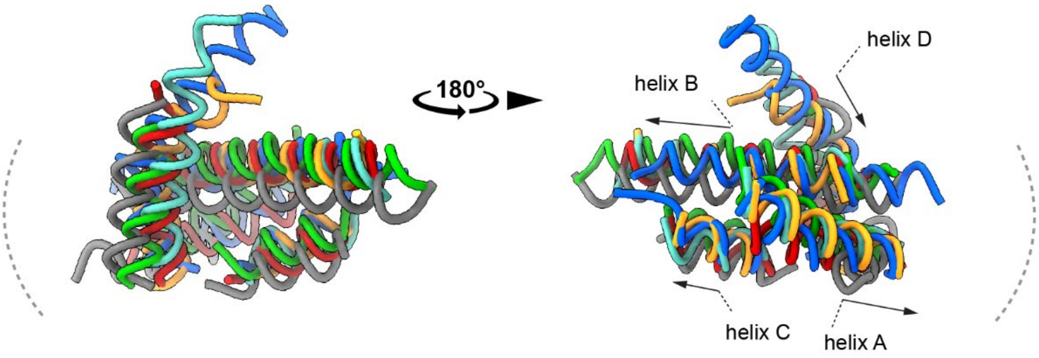


**Figure S12. Conservation of core helices including human picobirnavirus CP.**

Alignment of the core helices of representative CP structures from each recognised partitivirus genus and human picobirnavirus, annotated with the direction of the helix. The colour of the conserved helical core corresponds to the individual genus members shown in figures 4 and 5 of the manuscript: alphapartitivirus (AF2 - YP_086755) in red; betapartitivirus (PDB - 8PHH) in cyan; gammapartitivirus (PDB - 3ES5) in blue; cryspovirus (AF2 – YP_009508066) in yellow; deltapartitivirus (PDB - 7NCR) in green, picobirnavirus (6Z8D) in black.
